# A Programmable Model for Exploring the Functional Logic of the *Drosophila* Mushroom Body

**DOI:** 10.1101/2022.09.10.506218

**Authors:** Aurel A. Lazar, Mehmet Kerem Turkcan, Yiyin Zhou

**Affiliations:** Department of Electrical Engineering, Columbia University, New York, NY 10027, USA

## Abstract

We advance a model of the Mushroom Body of the *Drosophila* brain based on the recently published fruit fly brain connectome. We quantify the effect of the connectivity from antennal lobe projection neurons to Kenyon Cells (KCs) and the effect of feedback between KCs and the anterior paired lateral (APL) neuron on the representation of odorants at the KC level. We then characterize odorant representation in the mushroom body output neurons (MBONs) as a function of semantics represented by the dopaminergic neurons (DANs). Finally, we evaluate the performance of associative learning in the MB that endows pure odorants with DAN semantics.

## Executive Summary

### Overview

The MB circuit (Figure 1A) considered in this paper consists of 4 subsets of cell types: KCs, APL, MBONs and DANs. The connectivity of these cell types is specified by the 1) PN to KC connectivity (Figure 1B), 2) KC and APL connectivity (Figure 1C), 3) KC to MBON connectivity (Figure 1D), and 4) DAN to KC connectivity (Figure 1E).

**Figure 1:**
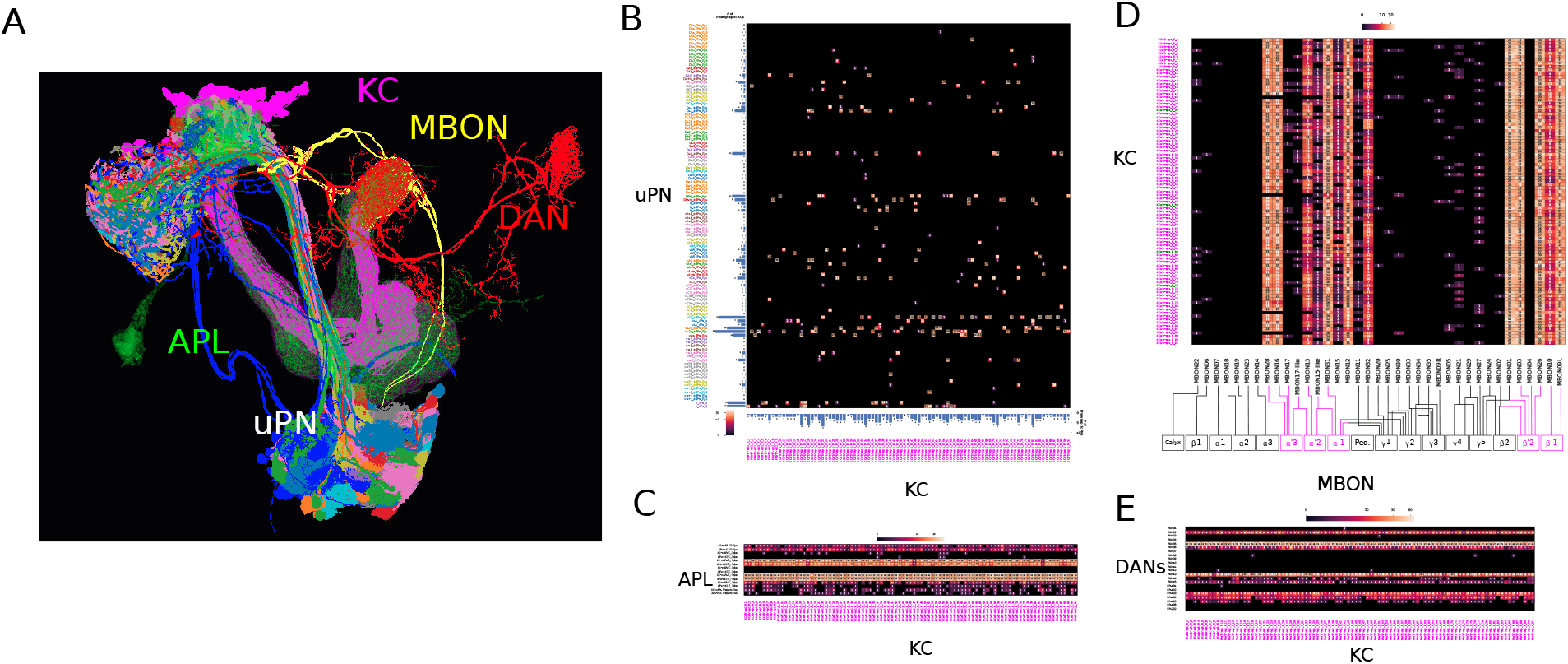
Cell types and cell connectivity in the Mushroom Body. (The figure only shows the connectivity of the *α′β*′-ap1 KCs; the connectivity of other KC types can be similarly constructed) **(A)** 4 Cell types of the MB. (magenta) KC (only *α′β′*-ap1 KCs shown). (green) APL. (yellow) MBON (only MBON14 shown), (red) DAN (only PPL106 shown). **(B)** Connectivity matrix between presynaptic uPNs and postsynaptic KCs (only *α*′*β*′-ap1 KCs shown). Bar chart on the x-axis shows the number of uPNs that provide inputs to a KC. Bar chart on the y-axis shows the number of postsynaptic KCs a uPN connects to. **(C)** Connectivity matrix between APL and KCs (only *α′β’*-ap1 KCs shown). The number of synapses from the APL to a KC and that from a KC to the APL are separately listed for the Calyx, the Pedunculus, *α*-, *β*-, *α*′-, *β*′- and *γ*-lobe. **(D)** Connectivity matrix between KCs and all 37 MBON types. (MBON09 on left hemisphere and MBON09 on the right hemisphere are treated here as two cell types as they arborize in different compartments in the MB of the left/right hemisphere). The number of synapses between a KC and all MBONs of a MBON type are aggregated and displayed as one entry in the connectivity matrix. MBON arborization of the lobes is listed under the x-axis, with the ones targeted by *α*′*β*′-ap1 KCs colored in magenta. **(E)** Connectivity matrix between 22 DAN cell types and KCs (only *α*′ *β*′-ap1 KCs shown). The numbers of synapses between DANs of the same cell type and a KC are aggregated and displayed as one entry in the connectivity matrix.

We model of the compartment circuit using the MB tensor that specifies the KC-to-MBON synapses that are modulated by DANs. In Figure 2A-F, we show the percentage of synapses of each KC-to-MBON connection that are modulated by the DANs. For example, in Figure 2A, the 100% (white patch) entry for KCa’b’-ap1_R_91 (last row) and MBON21 (column) indicates that the same entry in Figure 2D) is modulated by PAM02 DAN when the KC and the DAN are simultaneously active. Here, since the DAN-to-KC synapses are within 4*μm* of a KC-to-MBON synapse on the KC axon, the DAN is considered to be effective in modulating the KC-to-MBON synapse (the 4*μm* is somewhat arbitrarily picked).

**Figure 2:**
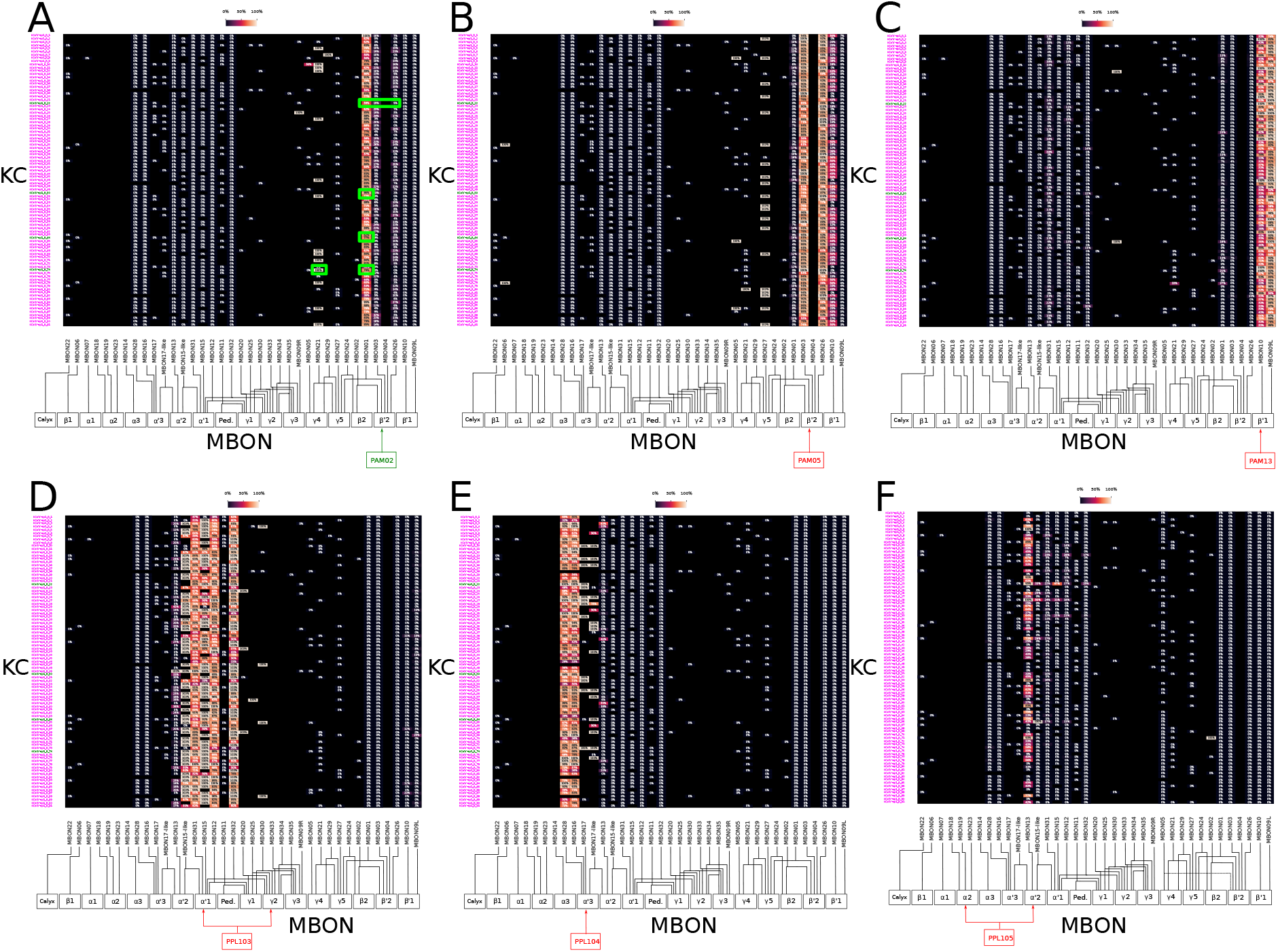
Neuromodulation of the KC/MBON connectivity by DANs. (The figure only shows the connectivity of the *α*′ *β*′-ap1 KCs; the connectivity of other KC types can be similarly constructed) **(A-F)** Percentage of each KC-to-MBON synapses that are modulated by a DAN cell type, as indicated by the green/red block at the bottom. (A) PAM02, (B) PAM05, (C) PAM13, (D) PPL103, (E) PPL104. (F) PPL105. Other DAN cell types are not shown. The diagrams at the bottom indicate the compartments that the DANs arborize. Here, a KC-to-MBON synapse is modulated by a DAN if any of the DAN-to-KC synapses is within 4*μm* of the KC-to-MBON synapse along the KC axon. Together, the matrices displayed in A-F, together with the one in Figure 1(D), form a 3-D tensor specifying the KC-MBON-DAN connectivity under neuromodulation.

The MB tensor is formed by stacking up the matrices in Figure 2(A-F) as well as those for all other DANs that are not shown. Assuming that patterns of associative memory reside in the KC-to-MBON connections/synapses, the MB tensor specifies which KC-to-MBON synapses are modulated by which DANs. Associative learning is achieved by modulating the entries of the MB tensor at the intersection between the odorant-activated KCs and the active DANs.

In Figure 2, for example, the KCs labeled in green are active when odorant A is present, and the DAN labeled in green is active. Consequently, the KC-to-MBON synapses in the MB tensor labeled in green are modulated.

### Representation of Odorants in the Mushroom Body

#### KC Representation of Odorants as a Function of the PN/KC Connectivity and APL Feedback

Here we quantify the KC odorant representation as a function of the PN/KC connectivity and APL feedback.

- Model PN-to-KC connectivity as a bipartite graph with a joint degree distribution.
- Model KC sparsification with APL feedback using connectome data.
- Evaluate the recoding with (110,110,3,15)-dimensional odorant-to-odorant representation tensor. We denote the representation tensor as **T**, with entries [**T**]_*o_1_,o_2_,m,c*_ where *o_1_* and *o_2_* denote 2 odorants, *m* ∈ [1, 3], and *c* ∈ [1, 15]. For a given compartment *c* and (i) *m* = 1, the tensor entry denotes the number of KCs activated by only odorant *o*_1_, (ii) *m* = 2, the tensor entry denotes the number of KCs only activated by odorant *o*_2_, and (iii) *m* = 3, the tensor entry denotes the number of (overlap) KCs that are activated by both odorants (individually).
- Quantify a measure of the tensor overlap and, using this measure, evaluate/compare different models of PN-to-KC connectivity (blue box in Figure 3) and APL feedback strength (red box in Figure 3). Average Percentage of Overlaps measures the average number of KCs that are activated by both odorants, over the total number of odorant pairs:

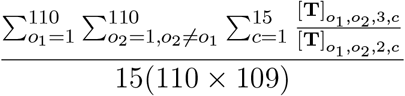

**Figure 3:**
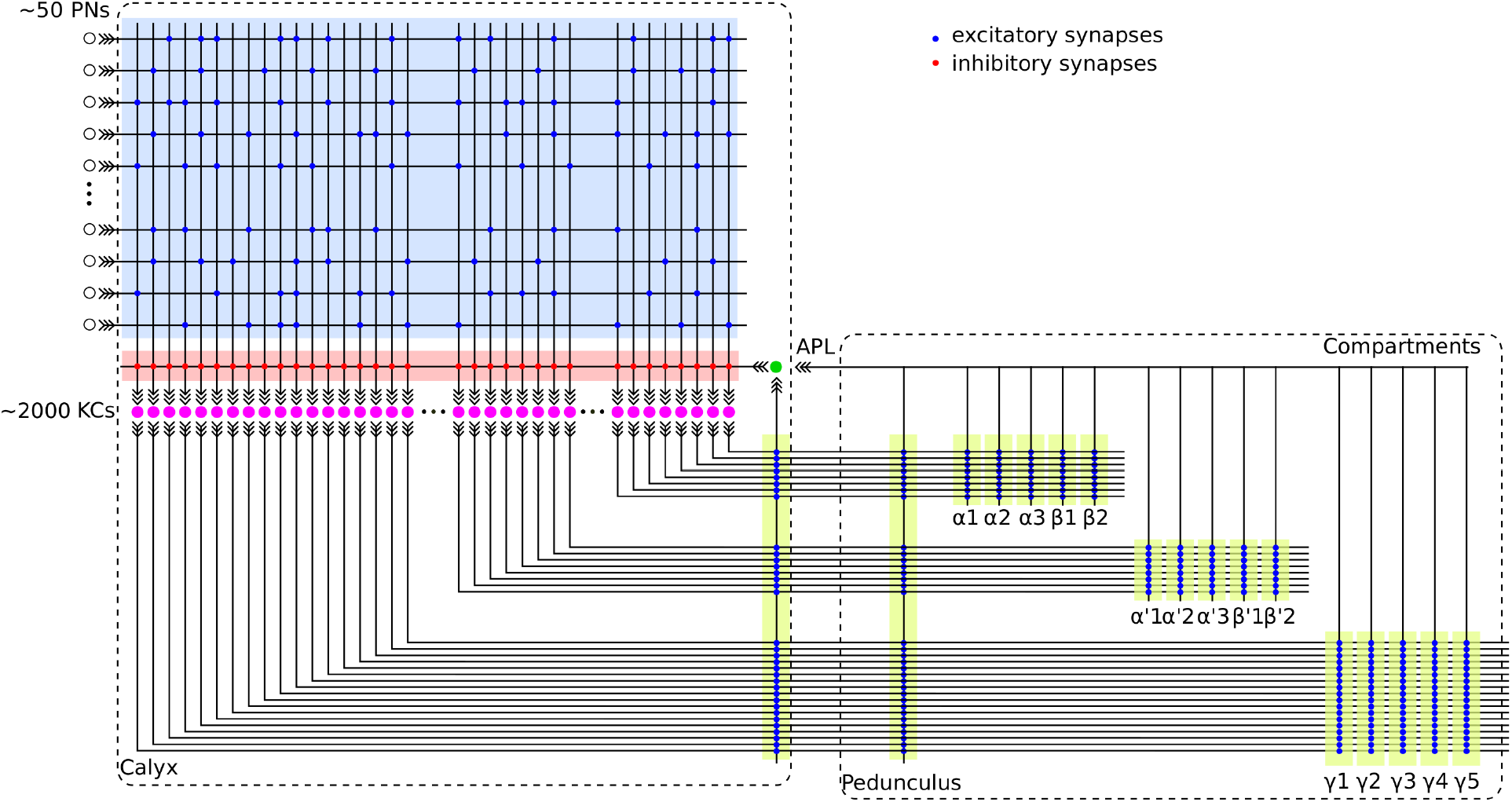
A circuit diagram of the PN-KC and KC-APL circuits.

The interpretation of this measure is that if this representation is passed onto the compartments, when odorant 1 is conditioned, what is the percentage of KCs (synapses) that are activate by odorant 2 will have already been depressed.

The effect of different models of PN-to-KC connectivity, the number of KCs and APL feedback strength (using number of feedback synapses) on this measure is shown in Figure 4.

**Figure 4:**
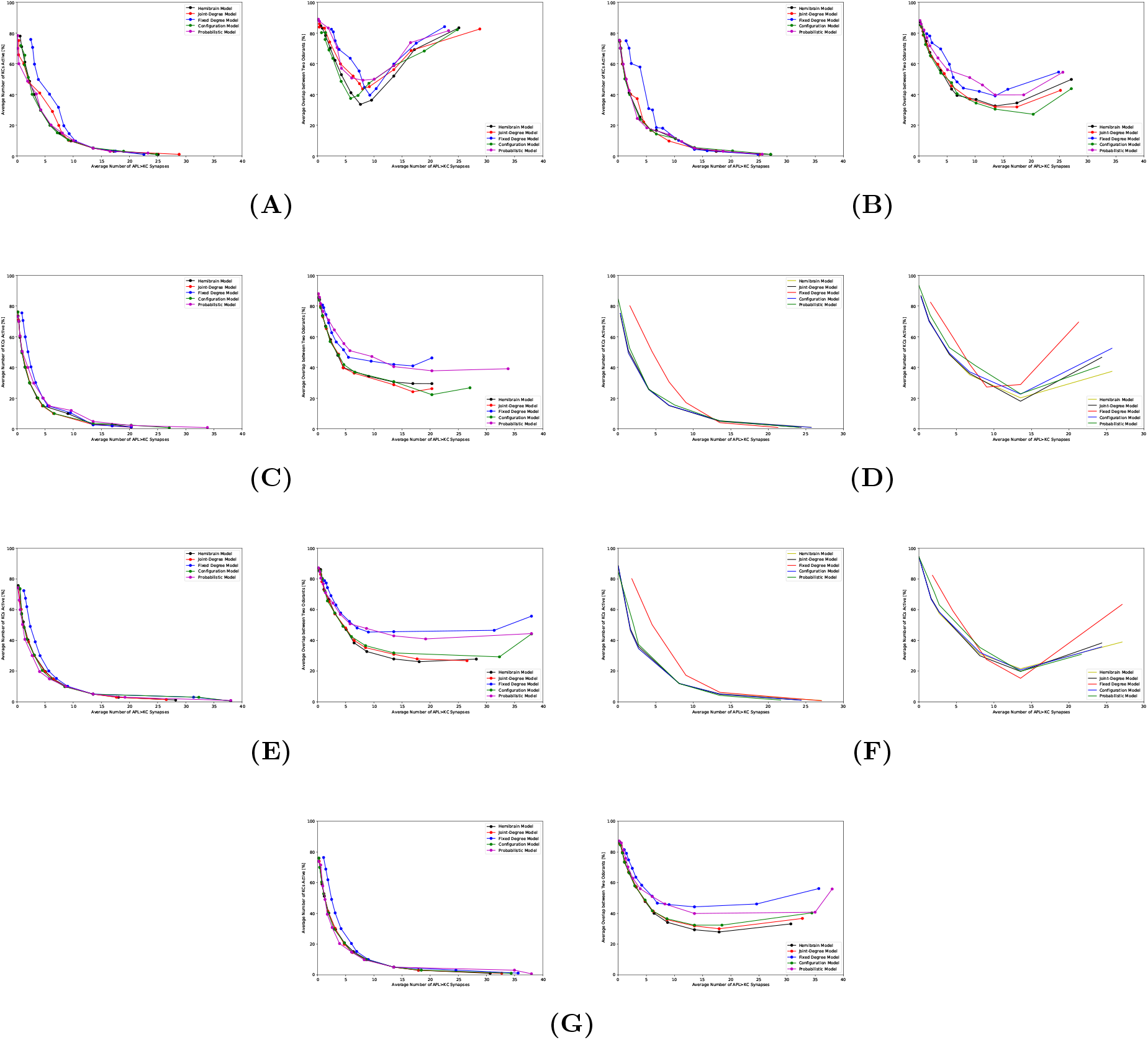
Average overlap percentage of the odorant representation tensor as a function of number of KCs, PN-KC bipartite graph model and average number of APL synapses onto KCs. **(A)** 10 KCs, **(B)** 100 KCs, **(C)** 400 KCs, **(D)** 800 KCs, **(E)** 1200 KCs, **(F)** 1600 KCs, **(G)** all 1472 KCs in Hemibrain data. (left) Average percentage of KCs activated by odorants. (right) Average overlap percentage.

### MBON Representation of Odorants as a Function of DAN Semantics

Here we quantify the MBON odor representation as a function of DAN semantics.

Determine from the vector of MBON responses which DAN inputs an odorant has been previously associated with.

- Assumption: 1) Every MBON will respond to any odorant input in the naive fly. 2) KC-to-MBON synapses are assumed to take the value 1 before modulation and 0 after modulation.
- Modeling the semantics of DAN: DAN semantics are specified in Figure 5 by the DANs that convey them into the compartments. Each US is represented by the activity of a vector of DANs.
- From MBON responses, we can tell if the presented odorant has been associated with specific DAN semantics. We track all MBON responses and if any of them is active, an odorant is present. The MBONs that are silenced indicate which of the DAN semantics the odorant has been associated with Figure 6.
- Evaluate the accuracy of associating odorant semantics and DAN semantics based on the connectome. For each DAN semantics, condition 1 out of the 110 odorants, evaluate the error rates when such procedure lead to a different odorant recalling the US semantics.
- Compare performance of different models of PN-to-KC connectivity and different APL feedback strength.

**Figure 5:**
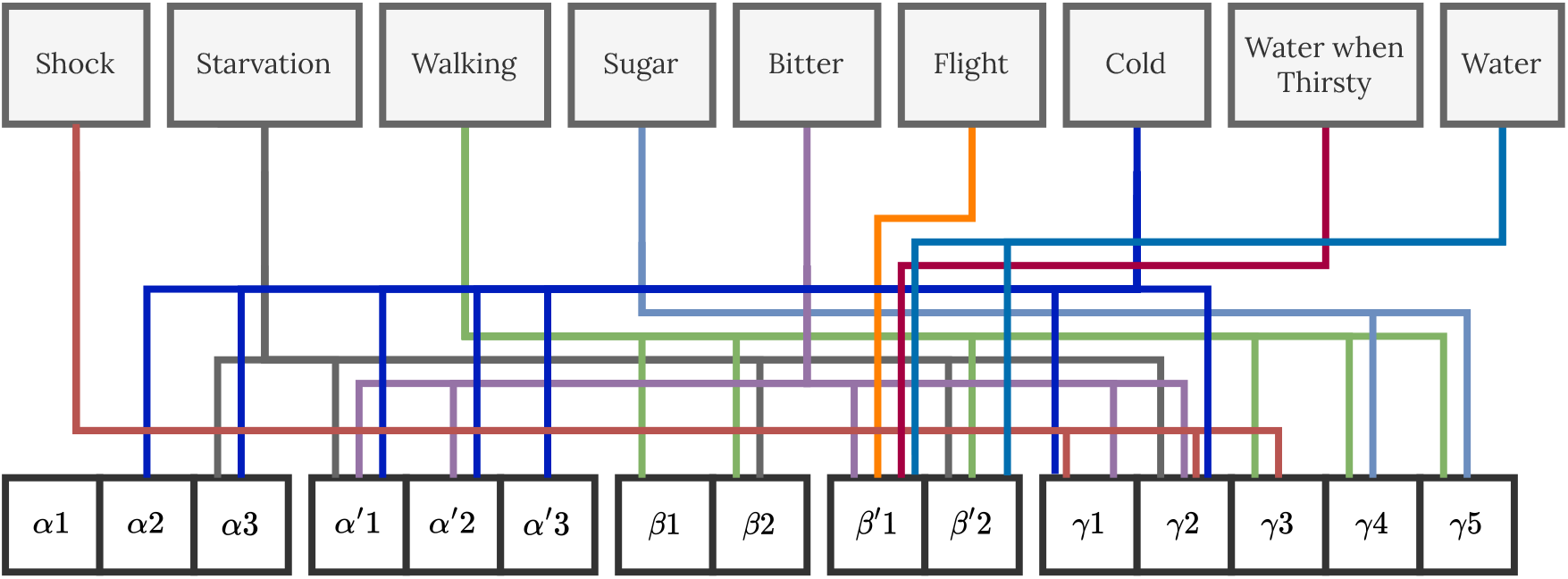
Semantic representation of DANs. We combine results from a wide variety of studies integrating various semantics: cold temperature [1], starvation [2], walking [3], sugar [4], electric shock [4, 5], bitter taste [3], flight [6], water [7].

**Figure 6:**
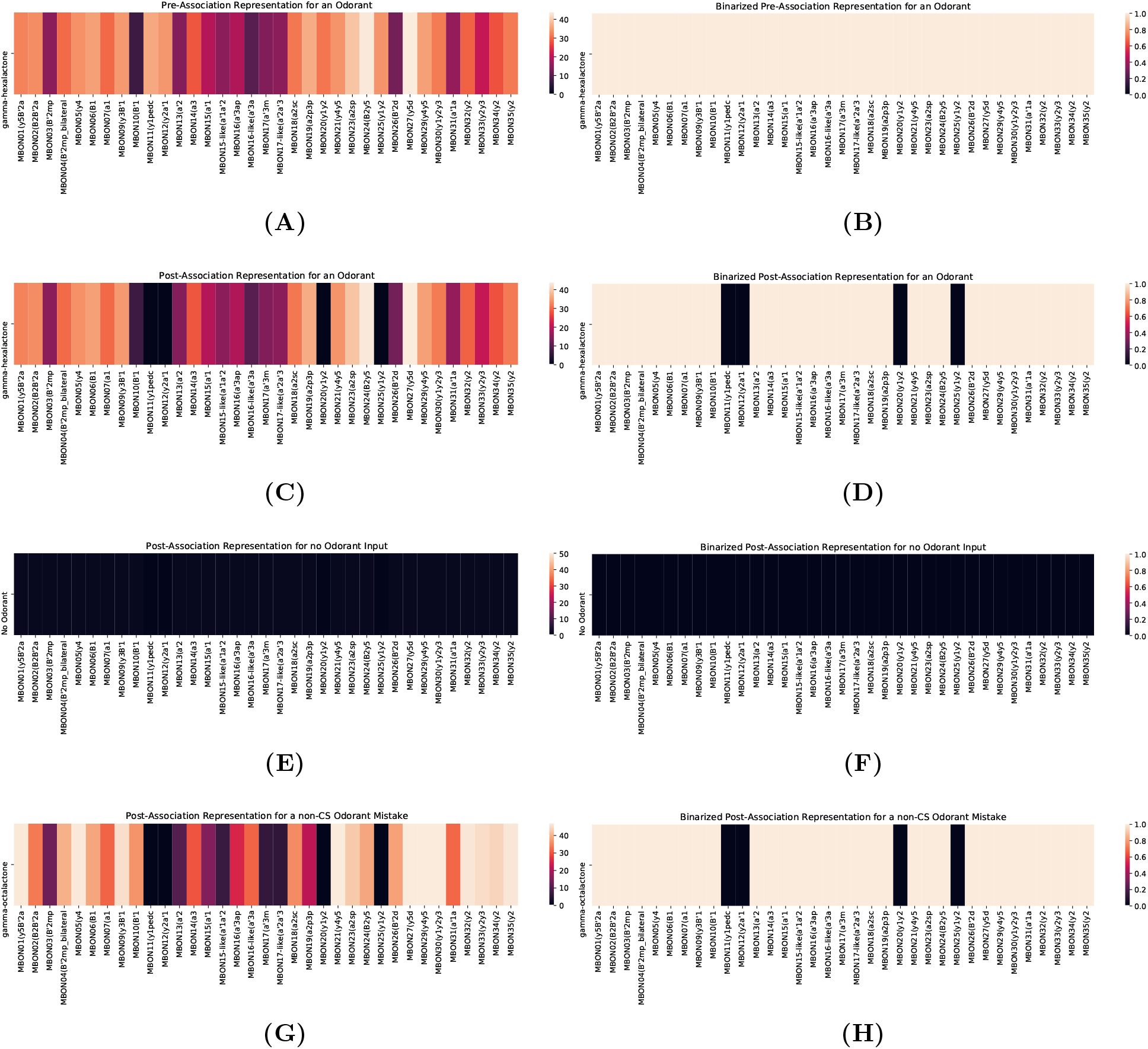
Visualization of the MBON odorant representation before and after an instance of associative learning. **(A)** The MBON response vector for an odorant. **(B)** Binarized response vector. **(C)** Post-association response vector when the odorant is used as the CS. **(D)** Binarized view of the odorant response vector. **(E)** When no odorant is present, there is no response across MBONs. **(F)** Binarized view of the odorant response vector cannot be used for the readout. **(G)** There can be mistakes in the MB readout. Here, a non-CS odorant gives no response in the US-related MBONs. **(H)** Binarized view of the odorant response vector similarly shows a readout mistake.

In Figure 7B, we associate 1 out of 110 odorants with electric shock and test if the changes in KC-to-MBON synapses caused by this association will lead to any of the other 109 odorants failing to activate all MBONs. If *N* of the other 109 odorant fail, then the error rate is *N*/109. We then repeat this procedure for all other 109 odorants. The overall error rate is the total error made divided by (110 × 109). The error rate is shown in Figure 7 evaluated for different models of PN-to-KC connectivity (color curves) and strengths of APL feedback (x axis). Each curve is U-shape because 1) when APL feedback is 0, the number of KCs activated by each odorant is high, leading to high overlap/error rate; 2) when APL feedback is too strong, KC representation is too sparse, and many odorants may not activate all MBONs even before the association occur, thus assumption 1) is not true any more and this leads to error.

**Figure 7:**
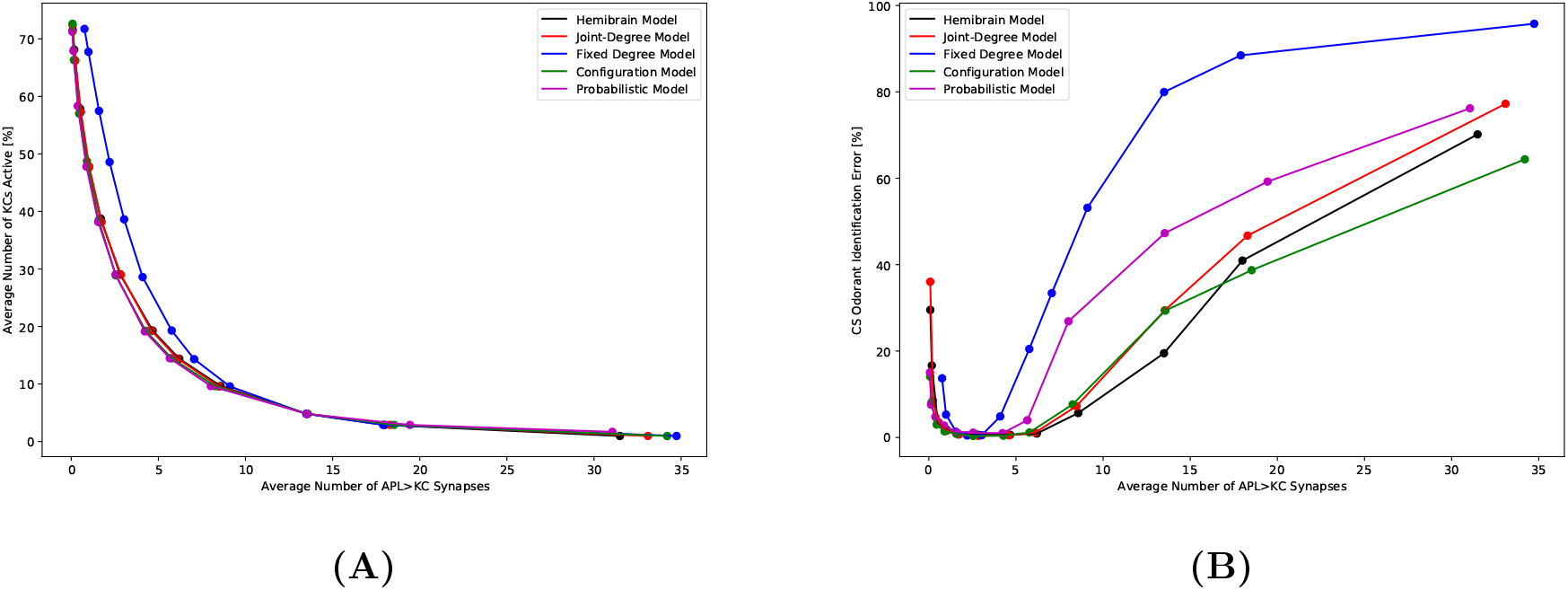
**(A)** Average power consumption vs KC-APL synapse strength (x axis) **(B)** Error rate vs KC-APL synapse strength (x axis) and different PN-to-KC connectivity models (color curves) for the electric shock modality. An error occurs when an odorant is mistakenly deemed, from the MBON response, as previously associated with a DAN semantics.

Figure 7 suggest a trade off between error rate of associative learning and the power consumption required for KC representation of odorants. The error rate is the lowest when about 50% of KCs, on average, respond to each odorant, and can be achieved regardless of which of the 5 uPN-KC circuit model is used. However, with uPN-KC circuit obtained from Hemibrain, joint-degree and configuration models, significantly reduction in the power consumption can be achieved when approximately 5-10% of KCs respond to each odorant, and without significantly impacting the error rate. For the other two models, we observe a rapid deterioration of associative memory performance below 15% KC sparseness.

## Acknowledgments

The research reported here was supported by AFOSR under grant #FA9550-16-1-0410, DARPA under contract #HR0011-19-9-0035 and NSF under grant #2024607.

